# Generation of clonal male and female mice through CRISPR/Cas9-mediated Y chromosome deletion in embryonic stem cells

**DOI:** 10.1101/2020.06.29.177741

**Authors:** Yiren Qin, Bokey Wong, Fuqiang Geng, Liangwen Zhong, Luis F. Parada, Duancheng Wen

**Affiliations:** Ronald O. Perelman and Claudia Cohen Center for Reproductive Medicine, Weill Cornell Medicine, New York, NY 10065, USA; Department of Medicine, Weill Cornell Medical College, 1300 York Avenue, New York, NY 10065, USA; Brain Tumor Center, Memorial Sloan Kettering Cancer Center, New York, NY 10065, USA

**Keywords:** Embryonic stem cells, CRISPR/Cas9, Y chromosome deletion, monosomic XO mice, clonal male and female mice, tetraploid complementation, genetically modified mouse models

## Abstract

Mice derived entirely from embryonic stem (ES) cells can be generated in one step through tetraploid complementation. Although XY male ES cell lines are commonly used in this system, occasionally, monosomic XO female all-ES mice are produced through spontaneous Y chromosome loss. Here, we describe an efficient method to obtain monosomic XO ES cells by CRISPR/Cas9-mediated deletion of the Y chromosome allowing generation of clonal male and female mice by tetraploid complementation. The monosomic XO female mice are viable and are able to produce normal male and female offspring. Direct generation of clonal male and female mice from the same mutant ES cells significantly accelerates the production of complex genetically modified mouse models by circumventing multiple rounds of outbreeding.

---

Genetically modified (GM) animals are essential tools for the study of both fundamental biology and human diseases. The production of GM animals relies on two critical features: 1) stable genome modifications and, 2) germline transmission of the mutations into a model system. A typical approach for creation of complex GM mice involves the generation of tetra-parental chimeras from normal embryos and GM embryonic stem (ES) cells, followed by multiple rounds of breeding to obtain both male and female homozygotes for germline propagation of the mutations. This process is time-consuming, laborious and costly, particularly if the final objective requires many independent germline manipulations in the same animal.

Mouse ES cells derived from the inner cell mass (ICM) of blastocysts have unlimited self-renewal and differentiation capacity if maintained in their ground-state pluripotency (1–3). Pure ES cell-derived mice (all-ES mice) can be directly and efficiently generated through tetraploid complementation, in which ground-state ES cells are injected into tetraploid blastocysts such that the host 4n cells can only contribute to the placenta but not somatic tissues (4–6). In this system by design, most viable animals are male, fertile female all-ES mice (39 chromosome, XO) are occasionally produced from the male ES cell lines (~2%) through spontaneous Y chromosome loss (7). Although the monosomic XO female (39, XO) mice have been proposed for the use of GM mice production to avoid mutant allele segregation during outcrossing (7), the observed low frequency makes it impractical for routine use in transgenic facilities. Here, we present a novel CRISPR/Cas9-mediated approach for directed elimination of the Y chromosome from mouse ES cells permitting efficient generation of monosomic XO female clonal mice by tetraploid complementation. The obtained monosomic XO female clonal mice are viable, fertile, and produce offspring of both sexes when crossed to male clonal mice from the same ES cells.

We derived new ES cell lines from hybrid F1 embryos by crossing C57BL/6j females with 129S1 males. The resultant male ES cell lines used in this study were all confirmed to produce normal all-ES mice by tetraploid complementation. Previous studies demonstrated that targeted chromosomal generation of multiple DNA double-strand breaks (DSBs) using CRISPR/Cas9 can induce directed chromosomal deletion (8, 9).

Thus, to eliminate the mouse Y chromosome, we targeted the RNA-binding motif gene *Rbmy1a1* which has over 50 copies exclusively clustered on the short arm of the Y chromosome (10) (**Fig. 1A**). We used synthetic gRNAs to target the *Rbmy1a1* gene sequences that have been successfully targeted to eliminate the Y chromosome in mouse embryos and ESCs (8). We used purified Cas9 proteins with nucleus localization signals (NLS) that can form functional gRNA-Cas9 ribonucleoprotein complexes (RNPs) in vitro. The use of pre-assembled gRNA-Cas9 RNPs allows for more accurate control of RNP composition and doses and has been shown to effectively reduce off-target effects and cytotoxicity in mammalian cells (11, 12).

**Figure 1.**
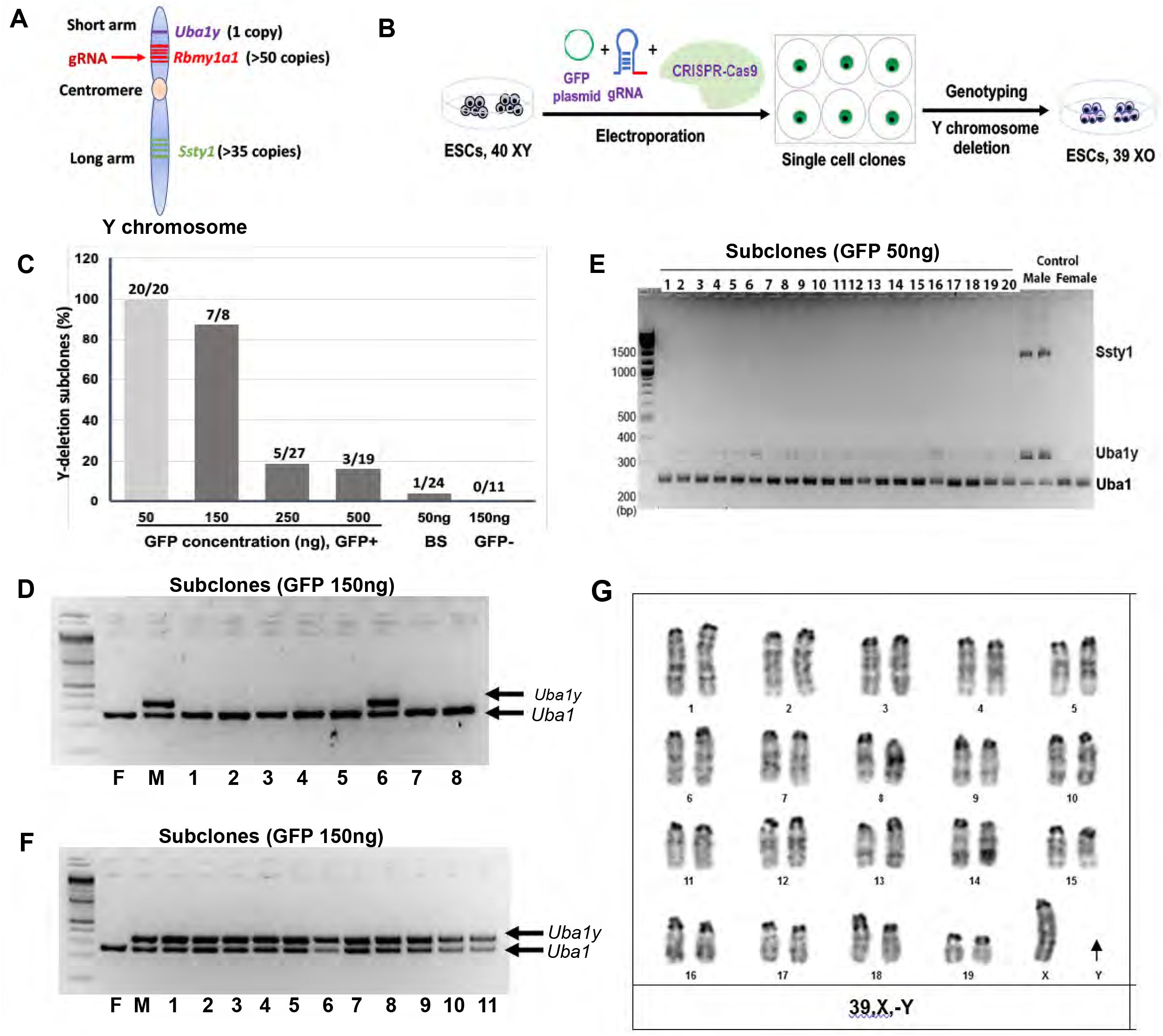
Efficient elimination of the Y chromosome in mouse ES cells using CRISPR/Cas9. A) Mouse Y chromosome and indicated relevant genes. B) Scheme illustration of CRISPR-mediated Y chromosome elimination in mouse ES cells. C) Y chromosome deletion rates as determined by *Uba1y* loss in targeted cells with varying concentrations of co-transfection GFP plasmid. BS: blinded selection. D) Genomic PCR analyses of *Uba1* and *Uba1y* genes in GFP^+^ subclones with 150ng GFP plasmid (36.9 nM). Female (F) and male (M) control DNAs were included. E) Genomic PCR analyses of *Uba1, Uba1y* and *Ssty1* genes in GFP^+^ subclones with 50ng GFP plasmid (12.3 nM). F) Genomic PCR analyses of *Uba1* and *Uba1y* genes in GFP^-^ subclones with 150ng of GFP plasmid. G) Karyotype analysis shows absence of the Y chromosome in a targeted ES cell (39, XO).

Electroporation is widely used to deliver RNPs due to the simplicity and large capacity (13). We included a circular GFP reporter plasmid with the Cas9 cocktail to allow for validation of successful electroporation. GFP expression could also serve as a surrogate, albeit indirect marker for Cas9-induced Y chromosome elimination. Indirect because plasmids and RNPs have differing modes of cell entry. Electroporation was performed using the Neon transfection system and continued cell culture after transfection. After ES cell colonies formed 48 hours post-electroporation, they were trypsin digested and single GFP+ cells were manually picked with the aid of a micromanipulator under a fluorescent microscope (**Fig. 1B and Supplementary Fig. 1A**). The GFP+ cells were individually plated in 96-well dishes for clonal expansion over feeder layers (**Fig.1B and Supplementary Fig.1A**). Cell colonies usually emerged after a week in culture and were expanded in gelatin-coated cultures for 1-2 passages before genotyping (**Supplementary Fig.1B**).

We first optimized our electroporation parameters by including a high dose of GFP plasmid DNA (123 nM, i.e., 500 ng/reaction) in the reaction along with gRNA-Cas9 RNPs at the supplier-recommended concentration (1.5 μM). Under the optimal electroporation condition, greater than 20% of the cells expressed GFP with minimal cell death. The GFP+ cell population did not vary in GFP plasmid concentrations ranging from 300-500ng although lower plasmid concentrations resulted in significantly decreased electroporation efficiency **(Supplementary Fig. 2C**).

We initially determined the state of the Y chromosome by genomic DNA PCR analyses for presence or loss of *Uba1y* gene located to the short arm on the Y chromosome and distal to the targeted *Rbmy1a1* gene (**Fig. 1A**). Three of nineteen clones showed loss of *Uba1y* gene in the 500ng electroporation group (**Fig. 1C and Supplementary Fig.1D**). Similarly, five of twenty-seven clones showed *Uba1y* gene loss in the 250ng plasmid electroporation (**Fig. 1C and Supplementary Fig. 1E**). Thus, the approximately 20% efficiency likely reflects conditions in which either: 1) CRISPR-induced chromosome loss occurs only in certain highly electroporation-receptive cells where excessive genomic breaks overwhelm cellular DNA repair capability or; 2) the excess uptake of GFP plasmids over Cas9 proteins could possibly generate a GFP+ cell insufficient Cas9 to induce enough DSBs for chromosome elimination, resulting in a GFP “blank-reporter”.

As Cas9 concentration is fixed in our system, we therefore next lowered the GFP plasmid concentration in the effort to mitigate the above concerns. Indeed, at the GFP plasmid concentration of 150ng, we found that seven of eight (87%) GFP+ subclones had lost the *Uba1y* gene (**Fig. 1C and 1D**), and at the lowest GFP plasmid concentration (50ng), 100% of the subclones (20 of 20) exhibited *Uba1y* gene loss (**Fig. 1C and 1E**).

We also examined the status of GFP-subclones in the conditions when essentially ~90% of GFP+ clones demonstrated efficient CRISPR/Cas9 targeting to assess whether the GFP reporter had value as a surrogate marker. While seven of eight GFP+ cell subclones yielded *Uba1y* gene loss, zero of eleven GFP-subclones had *Uba1y* gene loss in the 150ng group (**Fig. 1D and 1F**). In addition, of the twenty-four subclones derived from the cells blindly selected in the 50ng group, only one had *Uba1y* gene loss (**Fig. 1C**). These data confirm the values of the GFP plasmid in the appropriate ratio as a co-electroporation surrogate marker for successful gene transfer and likely successful gene targeting.

To further assess whether loss of the *Uba1y* gene effectively reflects deletion of the entire Y chromosome, we examined for retention of the *Ssty1* gene (>35 copies) that is located on the long arm of the Y chromosome **(Fig. 1A).** PCR analysis indicated the concomitant loss of the Y chromosome long arm gene, *Ssty1,* in each GFP+ subclone demonstrated to lack the *Uba1y* gene (**Fig. 1E**). Thus, loss of both *Uba1y* and *Ssty1* genes indicates that a large DNA segment, including the centromere of the Y chromosome, has been deleted. We further karyotyped the cell subclones with confirmed loss of both *Uba1y* and *Ssty1* genes using metaphase analyses. In the twenty metaphase chromosome spreads analyzed from clone-1, all cells displayed complete loss of Y chromosome with 18 cells having the expected 39 chromosome, XO karyotype (90%), while two cells were found with a segmental loss of 1q on chromosome 1 (**Fig. 1G, Supplementary Fig. 1E**). In a second clone (clone-2), all 22 cells examined showed XO karyotype with three cells showing additional abnormalities including an additional loss of 15q, chromosome 13 and an extra chromosome 14 (**Supplementary Fig. 1F**). These are likely *de novo* mutations arising during clonal cell expansion. In aggregate, these results confirm an efficient methodology for the physical elimination of Y chromosome by targeting double strand breaks in the *Rbmy1a1* gene in ES cells with minimal additional karyotypic perturbation.

The next critical step was to assess the potential and efficiency of CRISPR/Cas9 monosomic XO ES cells to generate all-ES mice by tetraploid complementation. ES cells with confirmed loss of the *Uba1y* and *Ssty1* genes were cultured and expanded for three to eight passages (total passages sixteen to twenty-one). We selected six of the twenty subclones (ES-1) with confirmed loss of the *Uba1y* gene (**Fig. 1E**) for blastocyst injection. Single cell clones from the parental ES cells without targeting were used as control. Of the six subclones tested, two clones gave rise to live pups with efficiency from 20% to 25% of the embryos transferred. A similar frequency (4 of 7 subclones) and efficiency (11%-21%) giving rise to all-ES mice were obtained for single cell clones of the parental ES cells (**Fig. 2A, Supplementary Table 1 and Table 2**). These data indicate that the preceding CRISPR/Cas9 manipulation of the ES cells did not adversely affect their pluripotency. In another test of four subclones generated from ES lines 2 and 3 (two subclones for each cell line) (**Fig. 1C**), three were able to generate live pups with similar efficiency to their parental ES cells by tetraploid complementation (**Fig. 2A**). As expected, pups obtained from the Y-deletion subclones were all of monosomic XO (39, XO) genotype and developed a morphologically normal female genital tract (**Fig. 2B and 2C**). Meanwhile, pups produced from the parental ES cells were all exclusively males (**Fig. 2C**). In agreement with other studies (7, 8), the monosomic XO female pups develop and mature normally to adulthood without noticeable defects (**Fig. 2C**). These results demonstrate the feasibility of efficient deletion of the Y chromosome from mouse ES cells using CRISPR/Cas9 technology allowing generation of male and female clonal mice from the same ES cell line.

**Figure 2.**
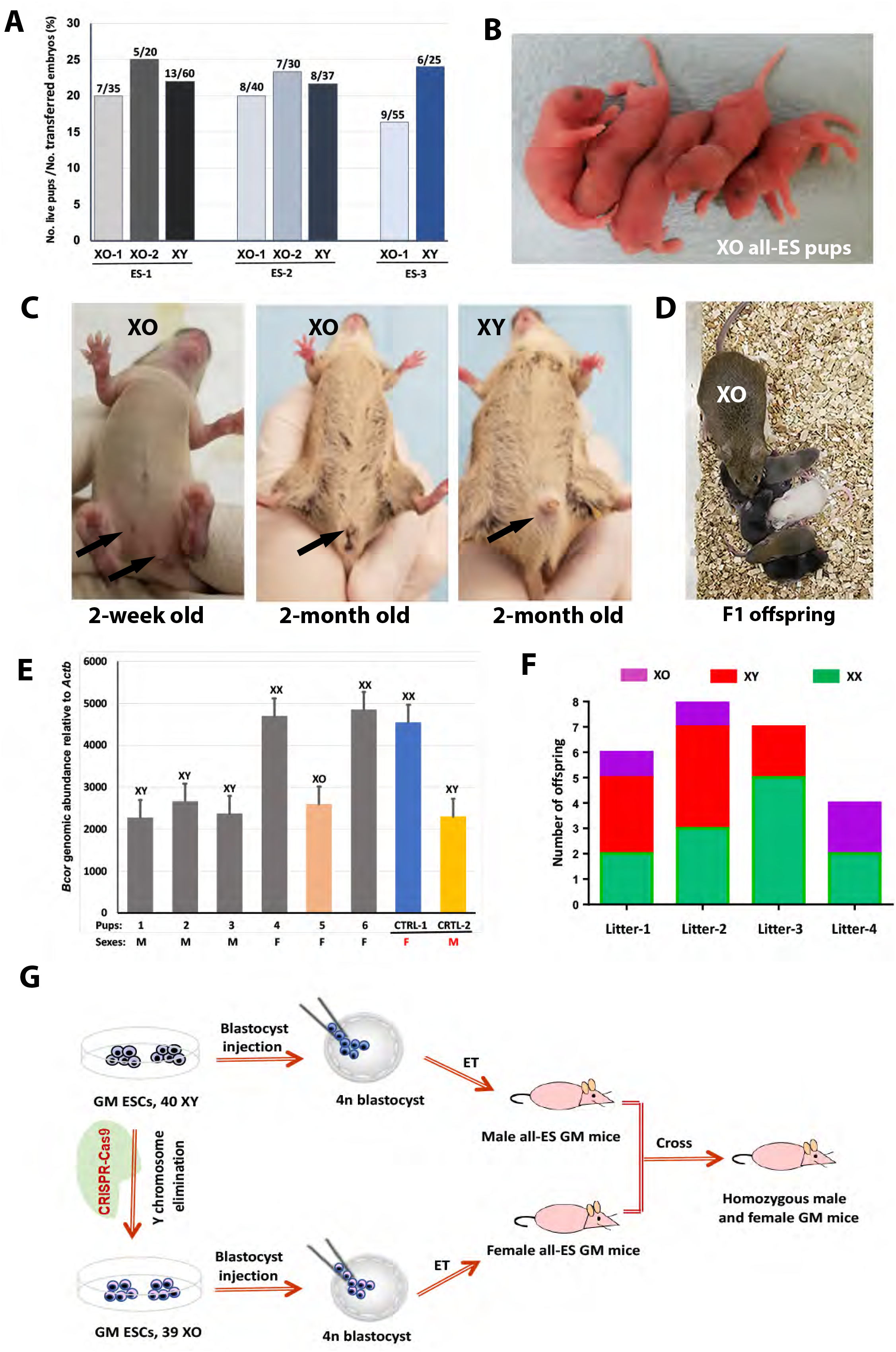
Viable and fertile monosomic XO female clonal mice generated from 39, XO ES cells. A) Developmental potential of Y chromosome deletion ES cells by tetraploid complementation. B) A litter of 5 XO all-ES pups from targeted subclone (ES-2). C) XO all-ES mice show a typical female genital tract at 2-week and 2-month, while all-ES mice from the parental ES cells show male genital tract morphology. D) XO female clonal mice produce normal offspring when mated with clonal male all-ES mice. E) QPCR quantification of genomic abundance of the X chromosome *Bcor* gene in indicated genomes of the offspring from litter-1. F) Normal male and female as well as XO offspring are produced by XO female clonal mice with a smaller litter size. G) A proposed approach to expedite the production of complex genetically-modified mouse models.

We further investigated monosomic XO female clonal mouse fertility of by breeding with clonal males from the same parental ES cell lines. All XO female clonal mice were fertile and delivered normal male and female offspring but with fewer littermates (4-8 pups) (**Fig. 2D–2F, Supplementary Table 3**). The XO mice generate two types of oocytes: X oocytes (19, X) and O oocytes (19, O), that following fertilization, result in four different genotypes (XX, XO, XY, OY). The X chromosome harbors essential developmental genes and therefore, OY embryos are expected to fail during embryonic development. To distinguish XO females from XX females in the offspring, we used qPCR to analyze copy number of an X chromosome single copy gene, *Bcor* (**Fig. 2E**). Monosomic XO females were also present among the offspring from the following generation with a frequency of one of three or four females (**Fig. 2F, Supplementary Table 3**). These numbers are lower than the expected Mendelian frequency of XO mice among females given that half of the females are expected to be XO. These results suggest that monosomic XO embryos may be less robust than XX embryos during development. Partial loss of XO embryos and embryonic lethality of OY embryos during the developmental stage would therefore account for the smaller litter size from XO mice. More importantly, the XO female clonal mice when bred with clonal males, give rise to both male and female offspring with normal genotypes (XY and XX) (**Fig. 2E–2F**). The production of normal XX and XY offspring from XO clonal mice indicates the feasibility to use XO all-ES mice for germline propagation of the complex genetic mutations in mouse models.

In conclusion, we demonstrate that targeting the Y chromosome single multi-copy *Rbmy1a1* gene in XY male ES cells using CRISPR/Cas9 technology can efficiently eliminate the Y chromosome. Importantly, the resultant 39 chromosome XO ES cells retain pluripotency and can generate viable and fertile all-ES mice that are phenotypically transgender from male to female. This system provides a practical strategy to manipulate sex in mice via ES cells, making it possible to expedite the production of complex multi-transgene GM mouse models which are a frequent necessity in current biomedical research, avoiding complex and time-consuming outcrossing (**Fig. 2G**).

## Methods

### Mice and embryos

Animals were housed and prepared according to the protocol approved by the IACUC of Weill Cornell Medical College (Protocol number: 2014-0061). Wild-type ICR mice were purchased from Taconic Farms (Germantown, NY). Females were superovulated at 6-8 weeks with 0.1 ml CARD HyperOva (Cosmo Bio Co., Cat. No. KYD-010-EX) and 5 IU hCG (Human chorionic gonadotrophin, Sigma-Aldrich) at intervals of 48 hours. The females were mated individually to males and checked for the presence of a vaginal plug the following morning. Plugged females were sacrificed at 1.5 days post hCG injection to collect 2-cell embryos. Embryos were flushed from the oviducts with KSOM+AA (Specialty Media) and subjected to electrofusion to induce tetraploidy. Fused embryos were moved to new KSOM+AA micro drops covered with mineral oil and cultured further in an incubator under 5% CO2 at 37°C until blastocyst stage for ES cell injection.

### Blastocyst injection

ES cells were trypsinized, resuspended in ES medium and kept on ice. A flat tip microinjection pipette was used for ES cells injection. ES cells were picked up in the end of the injection pipette and 10–15 of them were injected into each blastocyst. The injected blastocysts were kept in KSOM + AA until embryo transfer. Ten injected blastocysts were transferred into each uterine horn of 2.5 dpc pseudo-pregnant ICR females.

### ES cell line Derivation

ES cell lines were derived from hybrid F1 fertilized embryos of crossing between C57BL/6j females and 129S1 males. The ES cell derivation medium comprises 75 ml Knockout DMEM (SR, Gibco, Cat# 10829-018), 20 ml Knockout Serum Replacement (SR, Gibco, Cat# 10828), 1 ml penicillin/streptomycin (Specialty Media, Cat#TMS-AB-2C), 1 ml L-glutamine (Specialty Media, Cat# TMS-001-C), 1 ml Nonessential Amino Acids (Specialty Media, Cat #TMS-001-C), 1 ml Nucleosides for ES cells (Specialty Media, Cat# ES-008-D), 1 ml β-mercaptoethanol (Specialty Media, Cat# ES-007-E), 250 μl PD98059 (Promega product, Cat# V1191) and 20 μl recombinant mouse LIF (Chemicon International, Cat #ESG1107). The procedure to derive ES cell lines was described previously (4). Briefly, cell clumps originated from the blastocysts were trypsinized in 20 μl of 0.025% Trypsin and 0.75 mM EDTA (Specialty Media, Cat# SM-2004-C) for 5 min, and 200 μl of ES medium was added to each well to stop the reaction. Colony expansion of ES cells proceeded from 48-well plates to 6-well plates with feeder cells in ES medium, and then to gelatinized 25 cm^2^ flasks for routine culture in regular ES culture medium. Cell aliquots were cryopreserved using Cell Culture Freezing Medium (Specialty Media, Cat# ES-002-D) and stored in liquid nitrogen.

### CRISPR Cas9, gRNA, and ESC electroporation

Two crRNAs (IDT) targeting the *Rbmy1a1* gene at sequences 1) TTCAAGTGATGATGGTCTCCTGG and 2) TCCTTCATGTGAAGGGAACTTGG (including 3’ “NGG” PAM) (8) were annealed to a tracrRNA (IDT, cat# 1072533) at a 1:1:2 molar ratio to form dual duplex gRNAs by heating at 95 °C X 5 min and then cooled to room temperatures. Duplex gRNAs were then incubated with recombinant Cas9 protein (IDT, HiFi Cas9 nuclease V3, cat# 1081060) at room temperatures for 20 min to form gRNA-Cas9 ribonucleoproteins (RNPs), followed by co-delivery with a GFP plasmid (Addgene, #42028). All RNAs were in Duplex Buffer (30 mM HEPES, pH 7.5; 100 mM potassium acetate) and DNA in TE buffer (10mM Tris-HCl, 0.1mM EDTA, pH 7.5). The final concentration for each electroporation is 1.8 μM gRNA and 1.5 μM Cas9 nuclease.

To prepare ES cells for electroporation with the Neon transfection system (ThermoFisher), the cells were collected by trypsinization from culture, washed twice with PBS (without Ca2+ and Mg2+), and resuspended in supplied R buffer at 100,000 cells/10ul. 10 μl of cell suspension was mixed with 0.5 μl GFP plasmid DNA (27-270 mM) and 0.5 μl gRNA-Cas9 RNPs, and 10 μl of the mix was loaded to a Neon 10 μl tip for electroporation. Our optimized program is #14 (1200 volts, 20 seconds, 2 pulses). Treated cells were placed in a gelatin-coated 24-well with pre-warmed ES medium and returned to regular culture conditions.

### PCR Genotyping

PCR was performed on genomic DNA extracted from cell pellets or mouse tails to determine the loss of the Y chromosomal genes Uba1y and Ssty1 by utilizing KAPA Mouse Genotyping Kit (Roche, KK7302). Specific procedures were followed according to the reagent instructions. PCR products were separated on 2-3% agarose gel in TBA buffer, and inverted ethidium bromide stained images are shown. Cycling conditions: 95 °C for 3 min followed by 40 cycles of (95 °C for 15 s, 60°C for 15 s, and 72 °C for 120 s). Uba1/Uba1y primers: forward, TGGATGGTGTGGCCAATG; reverse, CACCTGCACGTTGCCCTT (335 bp product for Y-linked Uba1y, 253 bp for X-linked Uba1) (14). Ssty1 primers: forward, GCCACTATAGCTGGATTATGAG; reverse, GTCTTCACATCAGAGGTTCTAC (1,444 bp product) (8).

### qPCR of genomic DNA

To distinguish XX and XO female mice, X chromosome dosage was determined by measuring the abundance of the X chromosome resident *Bcor* gene relative to *Actb* in mouse tail genomic DNA using QPCR with PowerUp SYBR Green Master Mix (ThermoFisher, cat# A25778). Relative abundance of *Bcor* is presented as 1000*2^(Ct(Bcor)-Ct(Actb))^, where Ct is cycle threshold. Among several X-linked genes tested with this assay, only *Bcor* abundance in XX genome is consistently twice that in XY or karyotype-confirmed XO genome. *Bcor* primers: forward, TTTCCCACTCCATCCCCGACTAGTT; reverse, TCCCAAATAAACACCAGAGGCGACA). *Actb* primers: forward, GATATCGCTGCGCTGGTCGT; reverse, CCCACGATGGAGGGGAATACAG.

---

**Supplementary Table 1.** Developmental potential of Y-deletion subclones (ES-1) by tetraploid complementation

| Subclones | No. embryos transferred | No. live pups | %<br>(No. pups/No. embryos) |
| --- | --- | --- | --- |
| XO-1 | 35 | 7 | 20 |
| XO-2 | 20 | 5 | 25 |
| XO-3 | 20 | 0 | 0 |
| XO-4 | 20 | 0 | 0 |
| XO-5 | 20 | 0 | 0 |
| XO-6 | 20 | 0 | 0 |

**Supplementary Table 2.** Developmental potential of single cell subclones (ES-1) by tetraploid complementation

| Subclones | No. embryos transferred | No. live pups | %<br>(No. pups/No. embryos) |
| --- | --- | --- | --- |
| C1 | 57 | 0 | 0 |
| C2 | 17 | 2 | 11.8 |
| C3 | 53 | 7 | 13.2 |
| C4 | 60 | 13 | 21.7 |
| C5 | 43 | 0 | 0 |
| C6 | 61 | 9 | 14.5 |
| C7 | 20 | 0 | 0 |

**Supplementary Table 3.**
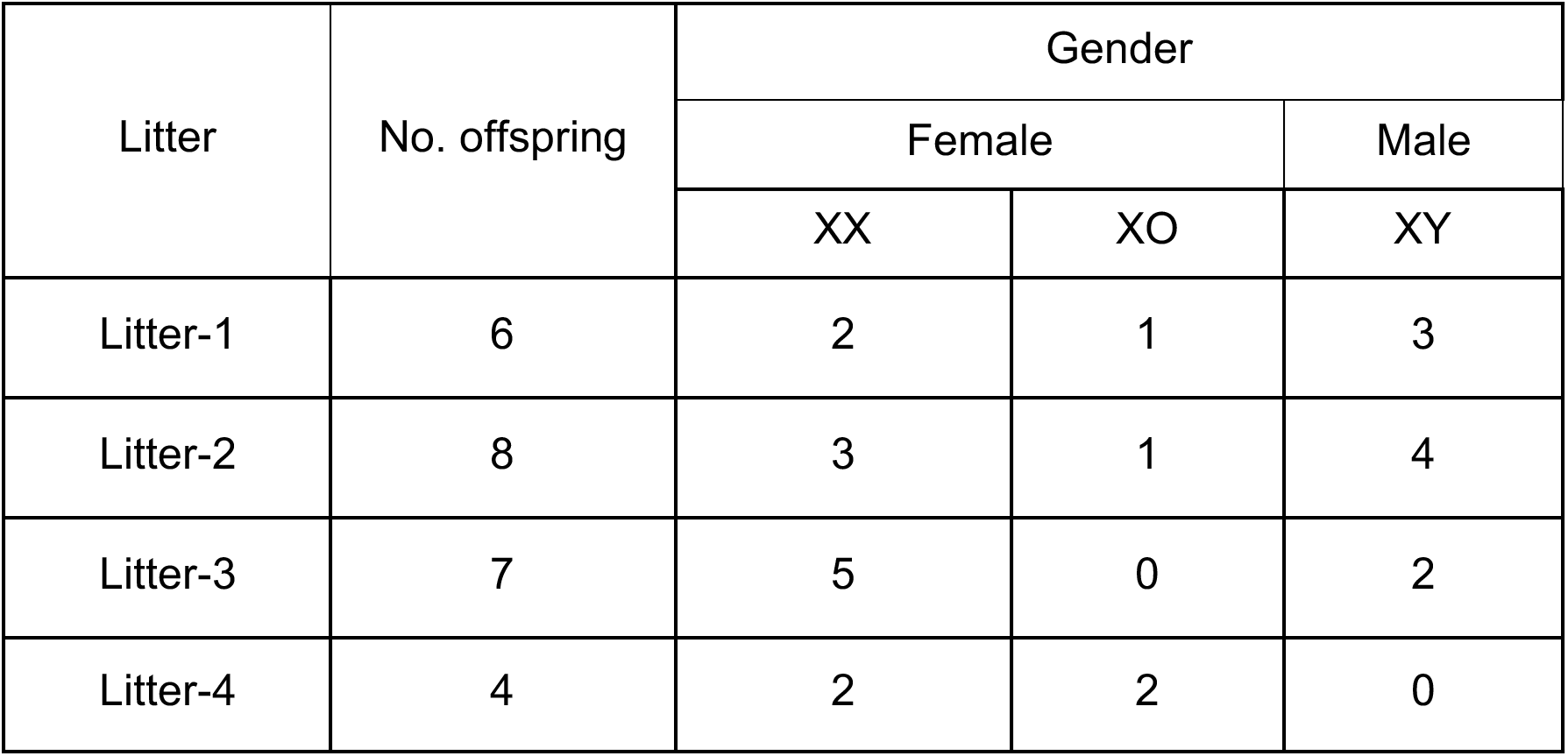
XO clonal females produce normal male and female offspring in cross with clonal males

| Litter | No. offspring | Gender |  |  |
| --- | --- | --- | --- | --- |
|  |  | Female |  | Male |
|  |  | XX | XO | XY |
| Litter-1 | 6 | 2 | 1 | 3 |
| Litter-2 | 8 | 3 | 1 | 4 |
| Litter-3 | 7 | 5 | 0 | 2 |
| Litter-4 | 4 | 2 | 2 | 0 |

**Supplementary Figure 1.**
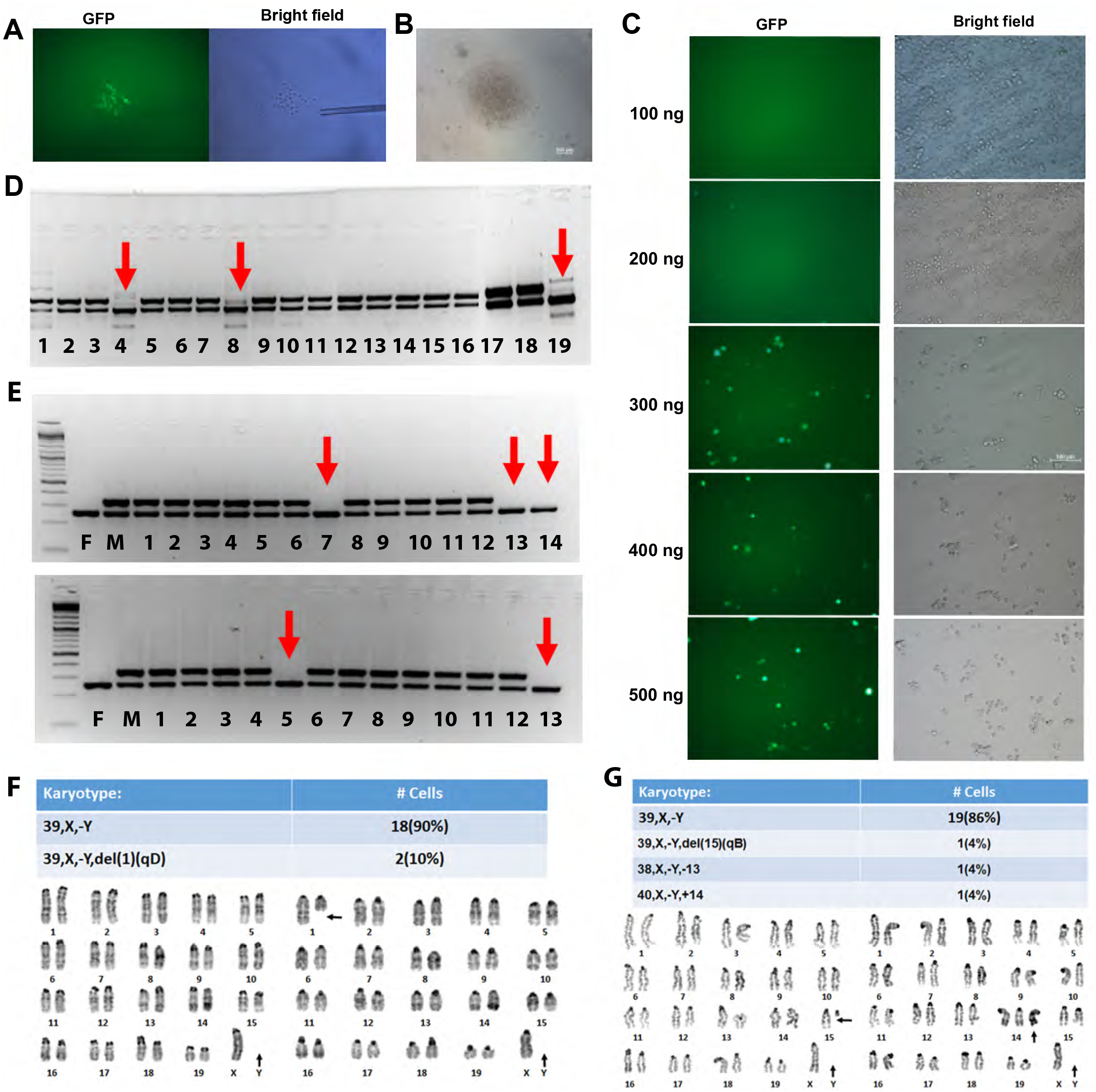
Efficient elimination of the Y chromosome in mouse ES cells using CRISPR/Cas9. A) GFP^+^ cells from the targeted ES cells were picked at 48h after transfection with the aid of a micromanipulator. B) A single cell colony from the GFP^+^ cells emerged on the feeders after one week of culture in ES medium. C) GFP^+^ cells with different GFP concentrations in the Cas9 cocktail. D) PCR genotyping of loss of *Uba1y* gene in indicated GFP^+^ subclones from the 500ng group. Red arrows indicate the *Uba1y* gene deletion clones. Male (M) and female (F) control DNAs were included. E) PCR genotyping of loss of *Uba1y* gene in indicated GFP^+^ subclones from the 250ng group. Red arrows indicate the *Uba1y* gene deletion clones. Male (M) and female (F) control DNAs were included. F) and G) Karyotyping of the subclones (ES-1) with loss of *Uba1y* and *Ssty1* genes confirms physical elimination of the Y chromosome.

## Notes

### Competing Interest Statement

The authors have declared no competing interest.

## References

1. Evans, M. J., and Kaufman, M. H. (1981) Establishment in Culture of Pluripotential Cells from Mouse Embryos. Nature 292, 154–156

2. Martin, G. R. (1981) Isolation of a Pluripotent Cell-Line from Early Mouse Embryos Cultured in Medium Conditioned by Teratocarcinoma Stem-Cells. P Natl Acad Sci- Biol 78, 7634–7638

3. Ying, Q. L., Wray, J., Nichols, J., Batlle-Morera, L., Doble, B., Woodgett, J., Cohen, P., and Smith, A. (2008) The ground state of embryonic stem cell self-renewal. Nature 453, 519–U515

4. Wen, D., Saiz, N., Rosenwaks, Z., Hadjantonakis, A. K., and Rafii, S. (2014) Completely ES cell-derived mice produced by tetraploid complementation using inner cell mass (ICM) deficient blastocysts. PloS one 9, e94730

5. George, S. H. L., Gertsenstein, M., Vintersten, K., Korets-Smith, E., Murphy, J., Stevens, M. E., Haigh, J. J., and Nagy, A. (2007) Developmental and adult phenotyping directly from mutant embryonic stem cells. P Natl Acad Sci USA 104, 4455–4460

6. Eggan, K., Akutsu, H., Loring, J., Jackson-Grusby, L., Klemm, M., Rideout, W. M., Yanagimachi, R., and Jaenisch, R. (2001) Hybrid vigor, fetal overgrowth, and viability of mice derived by nuclear cloning and tetraploid embryo complementation. P Natl Acad Sci USA 98, 6209–6214

7. Eggan, K., Rode, A., Jentsch, I., Samuel, C., Hennek, T., Tintrup, H., Zevnik, B., Erwin, J., Loring, J., Jackson-Grusby, L., Speicher, M. R., Kuehn, R., and Jaenisch, R. (2002) Male and female mice derived from the same embryonic stem cell clone by tetraploid embryo complementation. Nat Biotechnol 20, 455–459

8. Zuo, E. W., Huo, X. N., Yao, X., Hu, X. D., Sun, Y. D., Yin, J. H., He, B. B., Wang, X., Shi, L. Y., Ping, J., Wei, Y., Ying, W. Q., Wei, W., Liu, W. J., Tang, C., Li, Y. X., Hu, J. Z., and Yang, H. (2017) CRISPR/Cas9-mediated targeted chromosome elimination. Genome Biol 18

9. Adikusuma, F., Williams, N., Grutzner, F., Hughes, J., and Thomas, P. (2017) Targeted Deletion of an Entire Chromosome Using CRISPR/Cas9. Mol Ther 25, 1736–1738

10. Mahadevaiah, S. K., Odorisio, T., Elliott, D. J., Rattigan, A., Szot, M., Laval, S. H., Washburn, L. L., McCarrey, J. R., Cattanach, B. M., Lovell-Badge, R., and Burgoyne, P. S. (1998) Mouse homologues of the human AZF candidate gene RBM are expressed in spermatogonia and spermatids, and map to a Y chromosome deletion interval associated with a high incidence of sperm abnormalities. Human molecular genetics 7, 715–727

11. Kleinstiver, B. P., Pattanayak, V., Prew, M. S., Tsai, S. Q., Nguyen, N. T., and Joung, J. K. (2016) High-Fidelity CRISPR-Cas9 Nucleases with No Detectable Genome-Wide Off-Target Effects. Mol Ther 24, S288–S288

12. Kim, S., Kim, D., Cho, S. W., Kim, J., and Kim, J. S. (2014) Highly efficient RNA- guided genome editing in human cells via delivery of purified Cas9 ribonucleoproteins. Genome Res 24, 1012–1019

13. Kim, T. K., and Eberwine, J. H. (2010) Mammalian cell transfection: the present and the future. Anal Bioanal Chem 397, 3173–3178

14. Warr, N., Siggers, P., Bogani, D., Brixey, R., Pastorelli, L., Yates, L., Dean, C. H., Wells, S., Satoh, W., Shimono, A., and Greenfield, A. (2009) Sfrp1 and Sfrp2 are required for normal male sexual development in mice. Dev Biol 326, 273–284

